# An indicator cell assay for blood-based diagnostics

**DOI:** 10.1101/113357

**Authors:** Samuel A. Danziger, Leslie R. Miller, Karanbir Singh, G. Adam Whitney, Elaine R. Peskind, Ge Li, Robert J. Lipshutz, John D. Aitchison, Jennifer J. Smith

**Author notes:** To whom correspondence should be addressed: John D. Aitchison, Institute for Systems Biology, 401 Terry Ave N, Seattle, WA, 98109, Jennifer J. Smith Current address: PreCyte Inc. 401 Terry Ave N, Seattle, WA, 98109.

## Abstract

We have established proof of principle for the Indicator Cell Assay Platform™ (iCAP™), a broadly applicable tool for blood-based diagnostics that uses specifically-selected, standardized cells as biosensors, relying on their innate ability to integrate and respond to diverse signals present in patients’ blood. To develop an assay, indicator cells are exposed *in vitro* to serum from case or control subjects and their global differential response patterns are used to train reliable, cost-effective disease classifiers based on a small number of features. In a feasibility study, the iCAP detected pre-symptomatic disease in a murine model of amyotrophic lateral sclerosis (ALS) with 94% accuracy (p-Value=3.81E-6) and correctly identified samples from a murine Huntington’s disease model as non-carriers of ALS. In a preliminary human disease assay, the iCAP detected early stage Alzheimer’s disease with 72% cross-validated accuracy (p-Value=3.10E-3). For both assays, iCAP features were enriched for disease-related genes, supporting the assay’s relevance for disease research.

## Introduction

Prognostic and diagnostic blood biomarkers will cost-effectively and with limited patient risk, enable early detection, disease stratification, and assessment of response to treatment; but, their low abundance and the complexity of blood make their discovery challenging [1]. We have established proof of concept for the iCAP (Fig 1), a platform that can overcome these hurdles by using cultured cells as biosensors, circumventing the need to directly analyze molecules in serum by capitalizing on the natural ability of cells to detect and integrate weak biological signals in noisy environments. In the assay, patient serum or plasma is applied to cultured indicator cells and computational models identify disease state from the cells’ transcriptional response.

**Fig 1.**
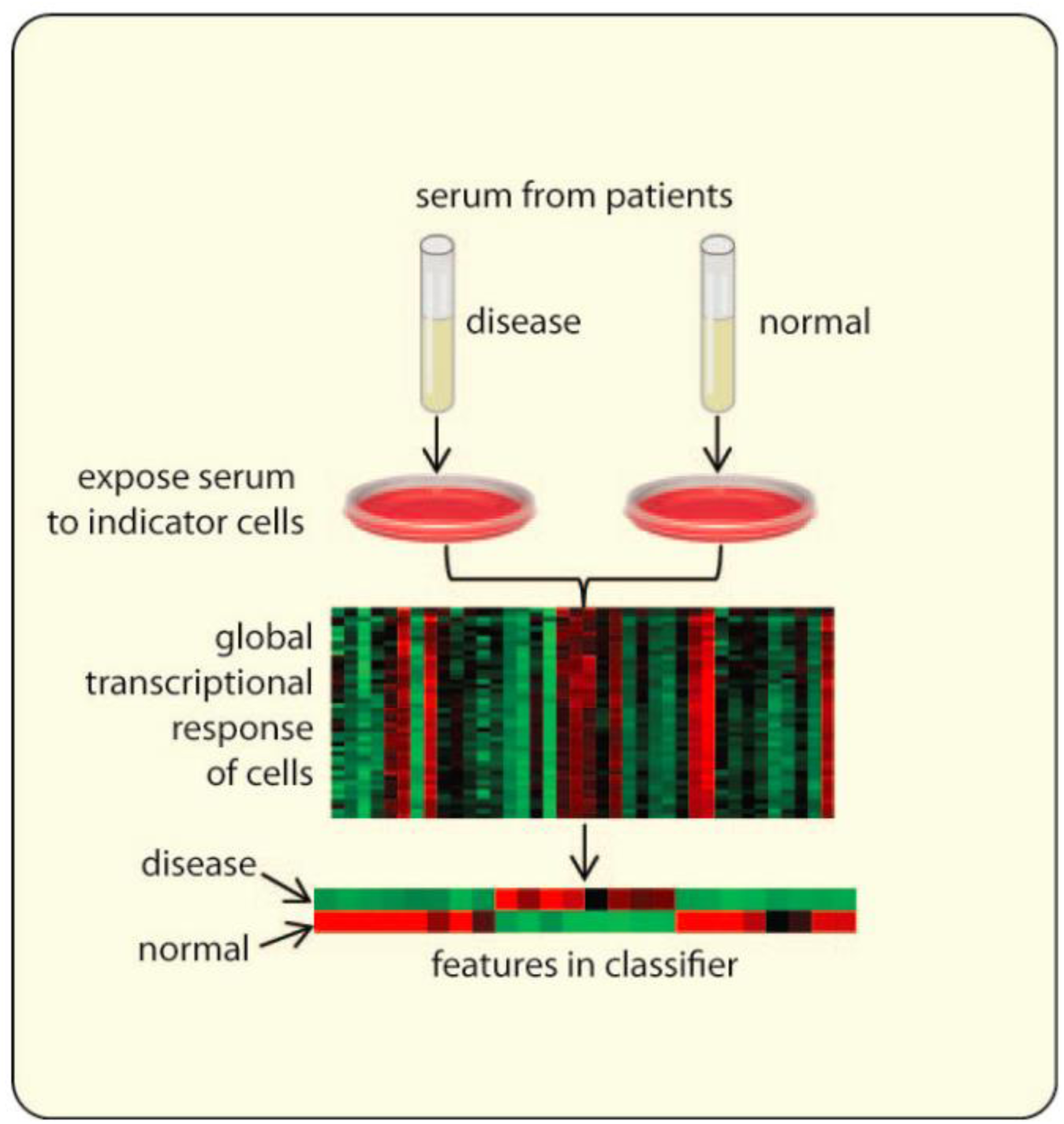
Development of an indicator cell assay. For each disease indication, the global differential gene expression pattern of the indicator cells is measured in response to serum from normal and diseased subjects, and is used to identify a reliable disease classifier using a small number of features. To deploy the assay, the cell based components can be automated and miniaturized, and the classifier genes can be measured using a simple, cost-effective and high-throughput readout in the same cell-based assay to classify new subjects.

Indicator cells are specifically selected for each application and generally are related to the disease being detected. To maximize clinical utility, assay development includes identifying robust standardized indicator cells that are genetically identical and reproducibly obtainable from stem cells. This approach with standardized cells was chosen over direct analysis of circulating cells from patients’ blood, to avoid noise arising from various aspects of individual patient cell heterogeneity.

The assay has several potential advantages over other diagnostic approaches, including: 1) Sensitivity — cells can detect blood components of low abundance, as even a single molecule can elicit a detectable cellular response [2]; 2) Specificity — the iCAP capitalizes on the naturally evolved ability of cells to detect specific signals in noisy environments, and field effects in which diseased cells can transform other cells nearby via secreted material [3,4]; and 3) Captures complexity — unlike most analyte detection devices, iCAP is not restricted to a single type of analyte because cells naturally respond to a broad range of molecules (including proteins, nucleic acids, metal ions, lipids and other metabolites, or their combinations).

The platform has potentially broad clinical applications for early diagnosis and monitoring progression for many diseases with signatures in blood. We present proof-of-concept of the iCAP for two different disease applications to demonstrate generalizability and versatility of the assay. First, we demonstrate an indicator cell assay for pre-symptomatic detection of disease in a mouse model of amyotrophic lateral sclerosis (ALS) with disease specificity when tested against samples from a Huntington’s disease mouse model. Second, we demonstrate an assay to detect Alzheimer’s disease (AD) at preclinical and early symptomatic stages from human plasma samples. These studies establish proof of concept for broad utility of the iCAP for early or preclinical detection of disease from blood samples. They support development of optimized and validated assays for clinical applications for both ALS and AD.

## Materials and methods

### Cell culture and gene expression analysis for ALS iCAP

An indicator cell assay was developed using previously frozen serum from a genetically engineered mouse model of ALS that reliably develops symptoms of disease at 80 +/− 10 days of age [5] (B6SJL-Tg(SOD1*G93A)1Gur/J expressing human mutant *SOD1* (G93A) from Jackson Laboratory (Main); colony 2726) [6]. Serum samples were from males that were hemizygous for the *SOD1* mutation (along with non-carrier littermates) at 65 +/− 6 days of age, an early stage of disease when subcellular unfolded protein response is detectable [7–9], but before symptom onset. (Note that the 12 serum samples used for independent testing of the classifier were all from mice at 64 days of age).

Indicator cells were embryoid bodies (EBs) containing spinal motor neurons, which were derived from mouse HGB3 embryonic stem cells (ESC) expressing a fluorescently labeled motor neuron marker (HB9-eGFP) (a kind gift of Hynek Wichterle and Thomas Jessell) using a published protocol [10]. Briefly, for each experiment, one vial of ES cells (~2 million cells) was plated in a 0.1% gelatinized T25 culture flask in ES cell medium (DMEM with 4500 mg glucose/L and 2250 mg Na-bicarbonate/L supplemented with 1× nonessential amino acids, 1× nucleosides (all from Millipore), 0.1 mM 2-mercaptoethanol (Sigma), 2 mM L-Glutamine (GIBCO), 1 × Penicillin/Streptomycin (GIBCO), 15% FBS (HyClone), and 1000 u LIF/mL (Millipore)). After 2 days (differentiation day 0), ES cells were trypsinized and replated at a density of 1–2 × 10^6^ cells per 10-cm dish (each in 10 mL of ADFNK (differentiation) medium) to induce embryoid body formation. ADFNK medium consisted of 1:1 Advanced DMEM/F12 (Life Technologies) and Neurobasal medium (Life Technologies) supplemented with 1X Penicillin-Steptomycin (Life Technologies), 200 mM L-glutamine (Life Technologies), 0.1 mM 2-mercaptoethanol (Sigma) and 10% knockout serum replacement (Life Technologies). On differentiation day 2, embryoid bodies were of fairly uniform size at 50-100-μm in diameter. One 10-cm dish of EBs was then split into 4 10-cm dishes in ADFNK medium. ADFNK medium was supplemented with 1 μΜ final concentration retinoic acid (RA) and 200 ng Hh-Ag1.3 agonist/mL (a kind gift of Curis, Inc.) for induction into spinal motor neurons. For a negative control, RA only was added to one dish. On differentiation day 6, fully induced EBs had bright signal from the HB9-eGFP reporter (by fluorescence microscopy), whereas signal from control EBs was not detectable. On day 6, efficiency of differentiation into fluorescently labeled motor neurons was estimated by dissociation of EBs into single cells using Accumax (Millipore) and counting percentage of fluorescent cells by FACS analysis. We measured ~30% of cells with detectable GFP fluorescence, which is consistent with the published efficiency of ~30–50% [10].

Six days after differentiation, EBs were plated in 12-well tissue culture plates at a density of ~0.25 million cells/cm^2^ in 2 mL of ADFNK medium. 5% mouse serum was added (either from *SOD1* mice or age-matched non-carrier controls) and incubated for 24 h. EBs were then harvested by centrifugation and RNA was isolated using Trizol^®^ according to the manufacturer’s recommendations. Total RNA samples were checked for quality (Agilent Bioanalyzer 2100) and quantity/concentration levels (NanoDrop ND-1000 spectrophotometer) and 0.1 μg of high quality RNA samples were processed for hybridization on GeneChip^®^ Mouse Exon ST microarrays (Affymetrix) following manufacturers recommendations to generate amplified and biotinylated sense-strand (ss) DNA targets from the expressed genome using Ambion^®^ Whole Transcript (WT) Expression Kit followed by GeneChip^®^ WT Terminal Labeling and Hybridization Kit (Affymetrix).

Gene expression microarray data from each batch of EBs were analyzed together using Affymetrix Expression Console™ software, and gene-level expression values were used to generate Pearson’s *r* scores for each array. Average Pearson’s *r* scores were evaluated using a Grubbs’ test (two-sided, alpha = 0.05) to identify outliers, which were removed from the data set.

### Developing an ALS classifier

Gene expression data was derived from exon array data using the Robust Multi-array Averaging (RMA) [11] algorithm as implemented in the [R] Aroma Affymetrix package [12]. Gene expression was estimated as the mean expression of all exons in each gene as annotated by release 33 of the Affymetrix MoEx-1_0-st-v1 annotation files. Next, gene expression levels were corrected for variations between batches of neurons. This correction was based on the levels on known motor neuron-related genes Olig2, Mnx1, Isl1, and Lhx3, chosen for their dynamic expression levels during motor neuron differentiation [10]. To do this, a linear model for each target gene was generated based on the expression of the four motor neuron genes in 12 assays with non-carrier serum samples from 5 different batches of neurons, and used as follows: GeneEx.log2 = log2((Gene Expression) / (Predicted Gene Expression)). Note that this model also served as a standard non-carrier gene expression profile reference for validation studies. Next, significantly differentially expressed genes and gene sets between carrier and non-carrier iCAP data were identified. For Gene Set Enrichment Analysis (GSA) [13], gene sets with GSA scores beyond a certain threshold (scores of −1 or +1) were selected.

To select differentially expressed genes and gene sets as features for disease classification, mProbes software [14] was used to rank gene or gene set expression features using 1000 tree deep Random Forest [15] feature selection. Those *N* ranked features that pass an mProbes FWER cutoff of 1.0 were used to build an ensemble of *N* classifiers. The first classifier in the ensemble used only the top ranked feature; the second classifier used the top two ranked features, and so on until the final classifier used all *N* features. Each classifier was allocated an unweighted vote and the probability of each class was estimated by the fraction of votes for that class across all classifiers.

For disease classification, first Minimum Entropy active learning was used to estimate the entropy of unlabeled examples by constructing a support vector machine [16,17] with a polynomial kernel and using a Laplacian distribution fitted on 3-fold cross-validated residuals to estimate class probabilities. Next, the single unlabeled example with the smallest entropy was selected to be predicted and labeled. Ties were broken using entropy as estimated using the ensemble predictions described previously.

### Study design for ALS iCAP

A sample size of 50 (25 of each class) for the classifier training set was determined informally as the number of samples expected to yield significant differential gene expression between the classes based on other studies in the literature. Because of the novelty of the approach, we did not have specific data sets appropriate for empirical sample size determination. Assays were performed in groups of 6 or 8, each with a different batch of differentiated motor neurons in embryoid bodies. Each batch was used to assay an equal number of randomly-selected disease and control samples. Before classification was attempted, the data from each batch of embryoid bodies were tested for outliers using Grubbs’ test (two-sided, alpha = 0.05) and a total of three outliers were removed. Classifiers were trained using the remaining 47 samples. The final configuration of Classifier 1 was locked down (on 12/10/14), and then blindly tested against 12 new unlabeled independent samples (on 12/11/14). Note that the 12 samples were not used for any prior classification or other analysis and the researcher performing the classification (S. Danziger) was blinded to the disease status of the samples, which were iteratively revealed during active learning by email. J. Smith and L. Miller each separately maintained the blinded data (in two different locations) and oversaw the evaluation by email communication after each round of active learning. The decision to test 12 independent samples was made prior to classifier testing and was based informally on the estimated accuracy of the classifier from cross-validation studies. Four additional classifiers were generated after this analysis and were tested using the same 12 independent samples used here, two of which are discussed in the Results section. Because five classifiers were generated, multiple comparisons were corrected for using Benjamini-Hochberg FDR control [18].

Classifier 2 and Classifier 3 used the same configuration as was locked down for Classifier 1 on 12/10/14 except for the specific modifications noted. While the reported accuracies are not truly blind as the labels for the validation set had been revealed to all researchers, they were performed to test specific aspects of the iCAP-based classification and were not chosen as high performing exemplars selected from a large number of additional classifier modifications.

Classifier 2 was trained to demonstrate classification with a small number of features and was tested on 01/12/15. Classifier 3 was trained to demonstrate classification with up- and down-regulated features for biological interest on 1/08/15. Two additional unnamed classifiers were tested on the 12-example validation set, but are not discussed in detail. One classifier (constructed on 1/9/15) used a GSA cutoff > 1 and, as expected from cross-validation studies, performed poorly (Accuracy = 0.67, p-Value = 7.30E-2, MCC = 0.35, p-Value = 2.59E-1, FDR [18] = 7.30E-2). The other classifier (constructed on 1/12/15) was a variant of classifier II that used a |GSA| cutoff > 1 (Accuracy = 0.75, p-Value = 1.93E-2, MCC = 0.58, p-Value = 4.93E-2, FDR [18] = 3.86E-2).

Note that no modifications, additions, or exclusions were made to the validation data set from the point at which the first classifier was locked down and neither the validation data nor any subset of it had ever been used to assess or refine the classifiers being tested. However, models were iteratively modified according the Active Learning method described and the unlabeled validation data was co-normalized with the training data as per the RMA algorithm. Neither of these actions violates the principal of a blind prediction since the classifier was never in any way exposed to the label of a validation example before that example was predicted.

The Huntington’s samples were known to all researchers to be samples from Huntington’s mice at the time of classification; however, the prediction methodology was a straightforward extension of the ALS classifier. The 12-example validation set was added to the 47-experiment training set to create a new 59 experiment training set. This data set was used to predict the class of the six Huntington’s samples using all 16 (i.e. 2^4^) logical combinations of the parameters ‘|GSA| ≥ 1 vs GSA ≤ −1’, ‘Gene Set vs Gene’, ‘mProbes vs no mProbes’, and ‘Active Learning vs no Active Learning’. These 16 sets of predictions were then repeated using 10,000 instead of 1,000 trees for the mProbes feature selection (S4 Table). Apart from the 6 Huntington’s samples, no other samples were predicted with any ALS classifier with the exception of two ALS samples that had undergone two freeze/thaw cycles. The effects of refreezing were inconclusive.

### Cell culture and gene expression analysis for AD iCAP

We developed an indicator cell assay to detect AD from human plasma samples at two stages of progression: preclinical (cognitively normal patients with positive CSF AD biomarkers; N=20) and early symptomatic (patients with mild cognitive impairment or early AD and positive CSF AD biomarkers) when compared to healthy controls (cognitively normal patients with negative CSF AD biomarkers). The study involved analysis of 18–20 samples of each class, which were processed in randomly selected batches, each containing an equal number of samples from each class. Indicator cells for this experiment were a commercially available, pan-neuronal population of glutamatergic and GABAergic neurons derived from induced pluripotent stem cells (iPSCs) (iCell Neurons from Cellular Dynamics International (CDI)). For this experiment, the neurons were thawed and plated in a 12-well dish (at 600,000 cells/well) and after culturing for 5 d, cells were exposed to 10% plasma containing 0.2 mg/mL heparin sodium salt in PBS (Stem Cell Technologies) for 24 h. RNA was isolated and used for gene expression analysis with Affymetrix human exon arrays (ST 1.0). Neurons were guaranteed free from mycoplasma contamination from CDI and were not tested again during the experiment.

### Developing the AD classifiers

Exon array data were analyzed using the Robust Multi-array Averaging (RMA)[11] algorithm as implemented in the [R] Aroma Affymetrix package [12]. Gene expression was estimated as the mean expression of all exons in each gene as annotated by release 33.1 of the Affymetrix human annotation files. The data for all samples were merged and normalized and analyzed for significant differential expression of genes and gene sets [13] between the classes as we have done for the ALS iCAP. Comparison of these data revealed significant overlap between the preclinical and early stage AD signatures. Therefore, the two disease classes were merged into one AD class (N=38) for comparison to the normal class (N=20). This approach had the advantage of increasing the disease sample size, increasing the power of the study.

Two different approaches were used to select features for constructing the classifier: the first selected genes as features, whereas the second identified *de novo* clusters of co-regulated genes as features. Like the gene sets used for the ALS classifier, the clusters were used to capture complex relationships between genes while reducing the number of features. For the gene-based feature selection, genes were ranked based on a joint probability score equally weighting three metrics: 1) magnitude of differential expression (fold change ratio), 2) significance of differential expression (t-test p-Value), and 3) significance of differential gene set expression (top GSA score observed for that gene using GSA algorithm [13]). For feature selection based on co-regulation (i.e. clustering), groups of co-regulated genes across all conditions were identified by analyzing all expression data (without class labels) with the cMonkey 2 biclustering software [19]. For this anlaysis, gene expression values were first converted to log transformed gene expression ratios using average gene expression values of four no serum controls (i.e. expression profiles of neurons not exposed to human serum) as a baseline.

AD classifiers were generated by combining gene and cluster features and ranking them using mProbes FWERs [14] and Random Ferns [20] feature selection. Features were selected using Sturges’ algorithm [21] to bin the FWERs and remove features with FWERs occurring in the 1.0 bin. Classifiers were then constructed using an SVM with a polynomial basis function optimized using the caret package with 10-fold cross-validation to optimize the area under the ROC curve. The primary accuracy of the classifier was determined using leave-one-out cross-validation and statistical robustness was evaluated by running 100 iterations of 16-fold cross-validation. The genes selected for these 1600 training sets (before the mProbes feature selection) were evaluated to determine the biological relevance of the iCAP cellular response.

### Study design for AD iCAP

Existing plasma samples were obtained from University of Washington Alzheimer’s Disease Research Center and were previously collected from subjects who provided written informed consent as part of a study approved by the institutional review boards of the University of Washington. Plasma samples were used to develop a blood-based assay to distinguish AD patients at two stages of AD progression from healthy controls (Tables 3 and 4). Both preclinical AD and normal subjects were chosen from clinically normal subjects based on CSF amyloid-β (1–42) protein fragment (Abeta-42) levels; both groups had clinical dementia rating (CDR) scores of 0, but preclinical AD samples had Abeta-42 levels ≤ 192 pg/mL, whereas normal samples had Abeta-42 levels > 192 pg/mL. In addition, preclinical AD samples were in the lowest percentile, whereas normal samples were in the highest percentile (based on levels of other subjects in the same clinically normal class in the biorepository), in order to make two groups more distinguishable biologically. Early symptomatic subjects had clinical diagnosis of either MCI or early AD, had CDR scores from 0.5–1, had Abeta-42 levels ≤ 192 pg/mL, and were in the lowest percentile of CSF Abeta-42 levels for that patient class. The CSF Abeta-42 threshold for defining the classes was a previously established threshold for AD diagnosis [22]. The three disease classes were matched for average age and gender. CSF collection and analysis were as described previously [23].

**Table 3:** Description of patient classes for AD iCAP

| class | N<sup>d</sup> | description | mean Abeta-42 (pg/mL) | Abeta-42 range (pg/mL) | average age (y) | male: female | average CDR<sup>e</sup> |
| --- | --- | --- | --- | --- | --- | --- | --- |
| preclinical AD<sup>a</sup> | 20 | cognitively normal; decreased CSF Abeta-42<sup>b</sup> | 129 | 67-186 | 73 | 11:9 | 0 |
| early symptomatic<sup>a</sup> | 18 | clinical MCI or early AD; decreased CSF Abeta-42<sup>b</sup> | 139 | 45-189 | 72 | 9:9 | 0.53 |
| normal | 20 | cognitively normal; no decrease in CSF Abeta-42<sup>c</sup> | 598 | 478-785 | 72 | 11:9 | 0 |

<sup>a</sup> Two disease classes were merged into a single early AD class. <sup>b,c</sup> Patient samples have CSF Abeta-42 ≤ 192 pg/mL and > 192 pg/mL, respectively. b and c are ranked in the lowest and highest percentiles for CSF Abeta-42 levels in their respective classes in the biorepository. CSF amyloid-β (1-42) fragment is abbreviated as Abeta-42. <sup>d</sup>Number of samples used to develop the classifier. <sup>e</sup>Average Clinical Dementia Rating.

**Table 4:** Leave-one-out cross-validated Alzheimer’s Disease Classifier

| Classifier Dataset | Accuracy | p-Value | Sensitivity | Specificity | MCC | MCC p-Values | AUC | AUC p-Values |
| --- | --- | --- | --- | --- | --- | --- | --- | --- |
| iCAP | 0.724 | 1.53E-04 | 0.838 | 0.524 | 0.382 | 3.10E-03 | 0.640 | 2.90E-02 |
| APOE | 0.672 | 2.68E-03 | 0.973 | 0.143 | 0.220 | 9.75E-02 | 0.564 | 4.12E-01 |
| iCAP.APOE | 0.690 | 1.12E-03 | 0.757 | 0.571 | 0.328 | 1.19E-02 | 0.689 | 4.68E-03 |

N = 58 samples, 38 MCI and pre-MCI Disease samples and 20 Normal Samples. No multiple testing correction was performed. MCC, Matthews correlation coefficient; AUC, area under the ROC curve.

The study was initiated with 21 samples of each class, but for 5 samples, RNA was not analyzed due to technical difficulties. Sample numbers were limited by the budget, and were justified based on power analysis of iCAP data with early symptomatic stage AD plasma samples and cognitively normal subjects (N=4 per class). A *t*-test was performed without multiple testing correction of exon-level expression values and 2,537 exons were individually significantly differentially spliced (p-value < 0.05). Next, a power calculation was performed [24], which suggested that a significant differential response signature of ~1000 exons will be generated using data from 20 paired disease/normal experiments.

### Data and software availability

All data needed to evaluate the conclusions in the paper are present in the paper and/or the Supplementary Materials. Microarray data related to this paper are available from GEO under the accession number XXX. Computer codes used in this publication are available from GitHub under the accession number XXX.

## Results

### Pre-symptomatic detection of disease in a mouse model of ALS

An indicator cell assay was developed for pre-symptomatic detection of disease in a transgenic *SOD1* (G93A) mouse model of ALS [6]. Disease mice of this model are hemizygous carriers of a mutant human *SOD1* gene (G93A) which reliably leads to the development of disease symptoms at 80 +/− 10 days of age [5]. Serum was from males that were either hemizygous for the *SOD1* mutation or non-carrier littermates at 65 +/− 6 days of age, an early stage of disease presentation when subcellular unfolded protein response is detectable [7–9] but typically before the onset of physical symptoms. Due to the mating strategy to maintain the colony, the mice are F1 progeny of two different strains (C57BL/6 and SJL), and have genetic heterogeneity outside of the *SOD1* transgenic locus [6], thereby challenging the robustness of the assay with genetic diversity.

Motor neurons were chosen as indicator cells because they are selectively degraded by ALS and have known responses to extrinsic signals of the disease [25]. Specifically, we used embryoid bodies (EBs) containing spinal motor neurons, which were derived from mouse HGB3 embryonic stem cells (ESC) expressing a fluorescently labeled motor neuron marker (a gift of Hynek Wichterle and Thomas Jessell) using a published protocol [10]. For this proof-of-concept study, motor neurons in EBs were used instead of dissociating EBs into individual cells in effort to minimize manipulation of the cells to reduce noise in the assay. Note that while EBs derived in-house were used for this initial study due to availability, standardized cells, which are more appropriate for a clinical assay are now commercially available, such as highly pure human motor neurons derived from iPSCs (available from iXCells, San Diego, CA).

Indicator cells were exposed to serum samples from pre-symptomatic *SOD1* mice (N=25) and age-matched non-carrier littermates (N=25) for 24 h. Global differential transcriptional responses were measured using exon-level Affymetrix^®^ microarrays, outlier experiments were removed and the remaining 47 data sets were co-normalized [26]. Gene expression level variation between batches of neurons was corrected using an in-house linear model based on expression of known motor neuron genes (see Methods).

### ALS Classifier 1 based on expression of 64 gene sets

These iCAP data were used to construct a disease classifier using average differential expression of gene sets as features (Classifier 1; as shown in Fig 2). Gene sets are sets of genes from databases (including GO, KEGG and REACTOME) with systematically annotated involvement in the same cellular process or pathway. This approach captured complex relationships between genes with a relatively small number of features (64 gene sets), which reduced the chance of overfitting the classifier to the available data. To do this, we first selected gene sets that were both significantly down-regulated in iCAP data with carrier versus non-carrier serum [13] and had more predictive potential than artificially generated random features [14]. Next, these gene sets were used to train an ensemble of support vector machine (SVM) disease classifiers [27,28]. Leave-one-out cross-validation selected 55–79 gene sets as features and correctly predicted 81% of the samples (binomial p-Value = 2.77E-06, Matthews correlation coefficient (MCC) = 0.62, Fisher’s transformation p-Value [29] = 3.73E-06).

**Fig 2.**
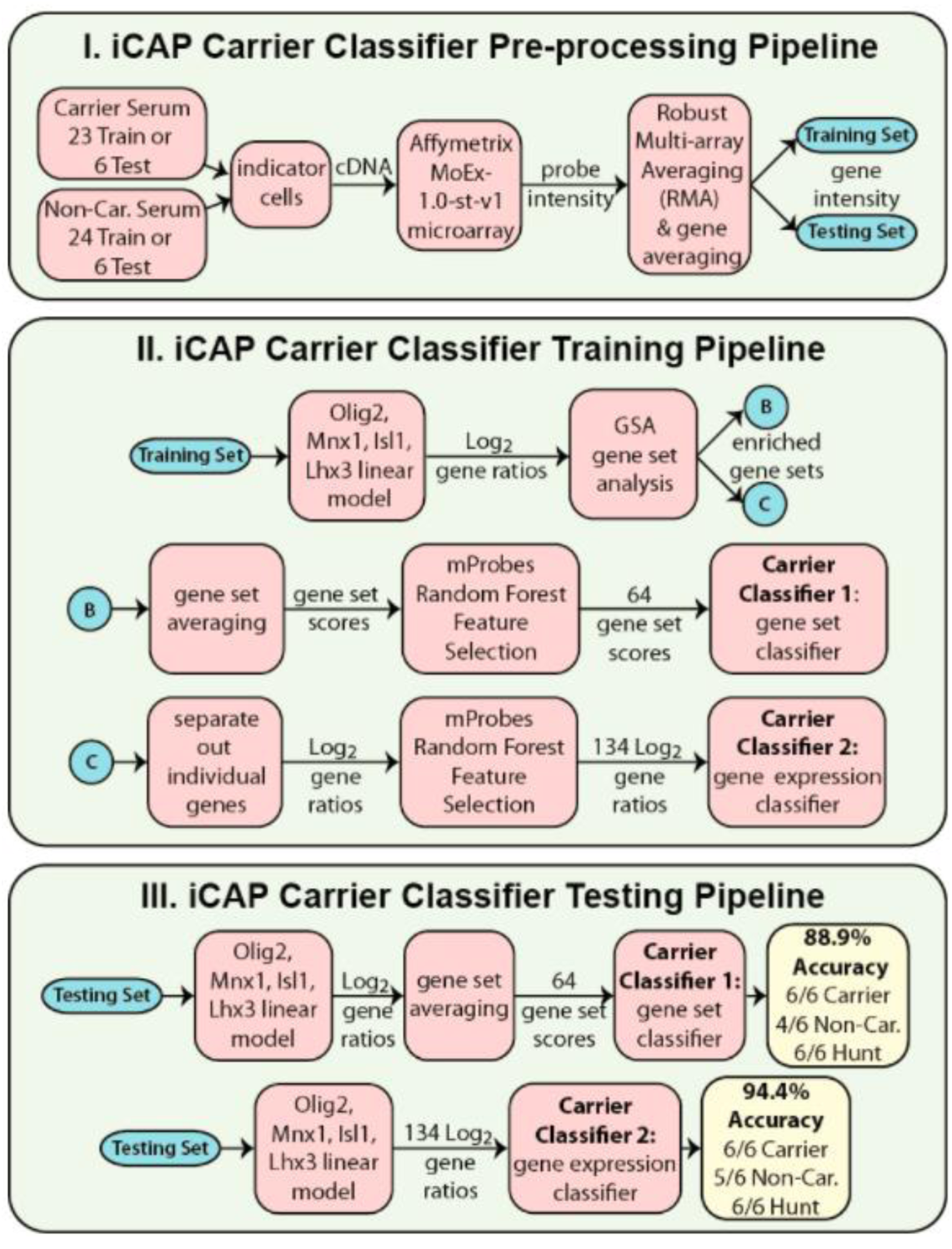
Classifier pipelines. I. 47 training samples or 12 test samples pass through the preprocessing pipeline, which results in normalized gene intensity scores. II. The classifiers are trained. 12 non-carrier (non-car.) samples in the 47 training examples are used to construct a linear model of gene expression based on control genes, which is used to convert all gene intensities into log2 gene expression ratios. These ratios are used to identify differentially expressed gene sets and ultimately select 64 gene sets or 134 genes that can differentiate between non-carrier and pre-symptomatic carrier mice using SVMs. III. The classifiers are tested. Gene intensities from 12 pre-processed but de-identified examples are converted to expression ratios using the linear model trained in II, and data sets composed of the 64 gene sets or 134 genes identified in II are extracted. The de-identified examples described by these data sets are classified as carrier or non-carrier using the SVMs trained in II. The pipeline for Classifier 3 is identical to that of Classifier 1 except that a different GSA score threshold was used (|GSA| ≥ 1 instead of GSA ≤ −1), resulting in selection of 106 gene sets instead of 64. For classifier testing with Huntington’s samples, classifiers were trained with all 59 normal and disease samples (not shown) and tested against 6 Huntington’s samples. For each classifier configuration, the fraction of Huntington’s samples that were correctly predicted as non-carriers of the ALS mutation is shown as ‘Hunt’ tally.

### Testing blind predictive accuracy of ALS classifier 1

To test the blind predictive accuracy of the ALS classifier, the classifier was first retrained using data from all 47 aforementioned serum samples, resulting in selection of 64 gene sets as features (S1 Table, Fig 2). Next, an independent test set was generated by assaying 12 new samples as before using serum from 6 pre-symptomatic *SOD1* mice and 6 non-carrier littermates, all at 64 days of age. For this test set, sample names were coded to hide the disease class of the mice. To test the classifier, we took advantage of a machine learning technique called ‘active learning’ [30] to make predictions on our blinded hold-out set one example at a time, revealing the disease class of that single example, and then retraining the classifier with the new data before we make a prediction on the next sample. This technique is appropriate for our proof-of-concept study because our dataset with ~20,000 measurements, but only 47 examples, is inherently under-determined. Active learning maximizes the size of the training set, allows the classifier to select the most robust models, and implicitly corrects for batch effects without violating the principle of blinded predictions. The classifier correctly predicted 6 of 6 pre-symptomatic *SOD1* mice and 4 of 6 non-carrier control mice, demonstrating 83.3% blind predictive accuracy (binomial p-Value = 3.17E-3; Table 1).

**Table 1:** Blind accuracies of ALS Classifiers

| actual class of<br>blind sample |  | predicted class of blind sample |  |  |
| --- | --- | --- | --- | --- |
|  |  | Classifier 1<sup>a,d</sup> | Classifier 2<sup>b</sup> | Classifier 3<sup>c</sup> |
| 1 | Non-carrier | *Carrier* | Non carrier | Non-carrier |
| 2 | Non-carrier | Non-carrier | Non-carrier | Non-carrier |
| 3 | Non-carrier | Non-carrier | Non-carrier | Non-carrier |
| 4 | Carrier | Carrier | Carrier | *Non-carrier* |
| 5 | Carrier | Carrier | Carrier | Carrier |
| 6 | Carrier | Carrier | Carrier | Carrier |
| 7 | Non-carrier | Non-carrier | *Carrier* | *Carrier* |
| 8 | Non-carrier | *Carrier* | Non-carrier | Non-carrier |
| 9 | Non-carrier | Non-carrier | Non-carrier | Non-carrier |
| 10 | Carrier | Carrier | Carrier | Carrier |
| 11 | Carrier | Carrier | Carrier | Carrier |
| 12 | Carrier | Carrier | Carrier | Carrier |
| performance metrics |  | Classifier 1 | Classifier 2 | Classifier 3 |
| Accuracy |  | 0.83 | 0.92 | 0.83 |
| p-Value |  | 3.17E-03 | 2.44E-04 | 3.17E-03 |
| q-value<sup>e</sup> |  | 5.28E-03 | 1.22E-03 | 5.28E-03 |
| Sensitivity |  | 1 | 1 | 0.83 |
| Specificity |  | 0.66 | 0.83 | 0.83 |
| MCC<sup>f</sup> |  | 0.71 | 0.85 | 0.67 |
| MCC p-Value |  | 1.01E-02 | 5.37E-04 | 1.79E-02 |
| **AUC<sup>g</sup>** |  | **0.88** | **0.90** | **0.89** |
| AUC p-Value |  | 1.14E-02 | 2.71E-03 | 1.12E-02 |

### Rationale for design methodology of ALS Classifier 1

The machine learning configuration used for the ALS classifier was determined by extensively testing many data subsets using cross-validation and other techniques to 1) test classifier robustness across various parameters, and 2) identify configurations that maximized accuracy and minimized the number of features to reduce the likelihood of overfitting. Therefore, Classifier 1 used aggregate gene set scores to reduce the number of features, and exclusively used gene sets down-regulated in disease samples, a configuration that tended to outperform those with up- and down-regulated gene sets in preliminary studies. We expect that as iCAP sample sizes become larger, and overfitting becomes a smaller problem, the optimal configuration will change to reflect finer grain models that more accurately identify and describe disease. Two additional configurations were tested after the initial blind predictions, and are presented below.

### ALS Classifier 2 based on expression of 134 genes

The 64 gene sets of Classifier 1 correspond to 263 genes, but for clinical applications, it would be more cost-effective to measure an even smaller number of genes. Therefore, we trained a second classifier using the expression levels of only the subset of these 263 genes that individually showed evidence of differential expression in response to carrier versus non-carrier serum (Fig 2). This approach aimed to remove genes that were members of a transcriptionally responsive gene set, but were individually not responsive. Using only the expression levels of these 134 genes, we trained a new classifier and used it to predict the disease status of the 12 independent samples. This gene expression classifier (Classifier 2, S2 Table) correctly identified 6 of 6 pre-symptomatic carrier mice and 5 of 6 non-carrier control littermates, demonstrating an accuracy of 91.7% (binomial p-Value = 2.44E-4; FDR[18] = 1.22E-3; Table 1), while measuring only a small number of genes.

### ALS Classifier 3 based on expression of 106 gene sets

For biological interest, we also trained an ALS classifier using the same parameters of Classifier 1, but with both up- and down-regulated gene sets (Classifier 3). This classifier, based on the expression of 106 gene sets (S3 Table), correctly predicted 5 of 6 pre-symptomatic *SOD1* mice and 5 of 6 non-carrier mice, demonstrating 83.3% predictive accuracy (binomial p-Value = 3.17E-03; FDR[18] = 5.28E-3; Table 1).

### ALS iCAP has disease specificity

ALS shares some pathobiology with other neurodegenerative proteinopathies such as Huntington’s disease. To test the disease specificity of the assay, we classified 6 serum samples from a mouse model of Huntington’s disease drawn at a pre-symptomatic age similar to that of the ALS model. 16 different classifier configurations were used, including those described above (Table 1) and others logically implied by the methodology (S4 Table). In aggregate, these classifiers had ALS sensitivity and specificity scores of 84%, and classified 95% of the Huntington’s samples as non-carriers of the *SOD1* mutation (p-Value = 2.09E-50). Also, adding Huntington’s samples (as non-carriers of the ALS mutation) to the test sets of the three classifiers in Table 1, tended to improve classifier performance (Table 2).

**Table 2:** ALS classifier accuracies with 6 Huntington’s disease samples included in test set

| Metric | Classifier 1<sup>a,d</sup> | Classifier 2<sup>b</sup> | Classifier 3<sup>c</sup> |
| --- | --- | --- | --- |
| Accuracy | 0.89 | 0.94 | 0.83 |
| p-Value | 7.25E-05 | 3.81E-06 | 6.56E-04 |
| Sensitivity | 1 | 1 | 0.83 |
| Specificity | 0.83 | 0.92 | 0.83 |
| FPR<sup>e</sup> | 0.17 | 0.08 | 0.17 |
| FDR<sup>f</sup> | 0.25 | 0.14 | 0.29 |
| F<sub>1</sub><sup>g</sup> | 0.86 | 0.92 | 0.77 |
| MCC<sup>h</sup> | 0.79 | 0.89 | 0.64 |
| MCC p-Value | 9.42E-05 | 9.71E-07 | 3.87E-03 |
| AUC<sup>i</sup> | 0.94 | 0.95 | 0.92 |
| AUC p-Value | 4.22E-04 | 1.11E-04 | 2.13E-03 |

### Indicator cell assay data reflect known aspects of ALS

The genes responding in the iCAP reflect biology underlying ALS, further demonstrating disease specificity. ER stress response mediated by PERK is a pathological event in sporadic and many forms of familial ALS [31]. This response involves the regulation of genes involved in protein synthesis by transcription factors (TFs) ATF4 and CHOP [32], and is prominently represented in the classifier feature sets: 1) The gene sets of Classifiers 1 and 3 include ‘PERK regulates gene expression’, ‘ATF4 activates genes’, and many cellular pathways related to protein synthesis that include known targets of ATF4 and CHOP (S1 and S3 Tables). 2) The 134 genes in Classifier 2 are significantly enriched for those directly targeted by ATF4 and/or CHOP in response to ER stress [32] (hypergeometric distribution (HGD) p-Value = 3.26E-28) (Fig 3). These data demonstrate that while the classifier was trained using a *SOD1* ALS model, the cellular response includes pathways common to multiple forms of ALS.

**Fig 3.**
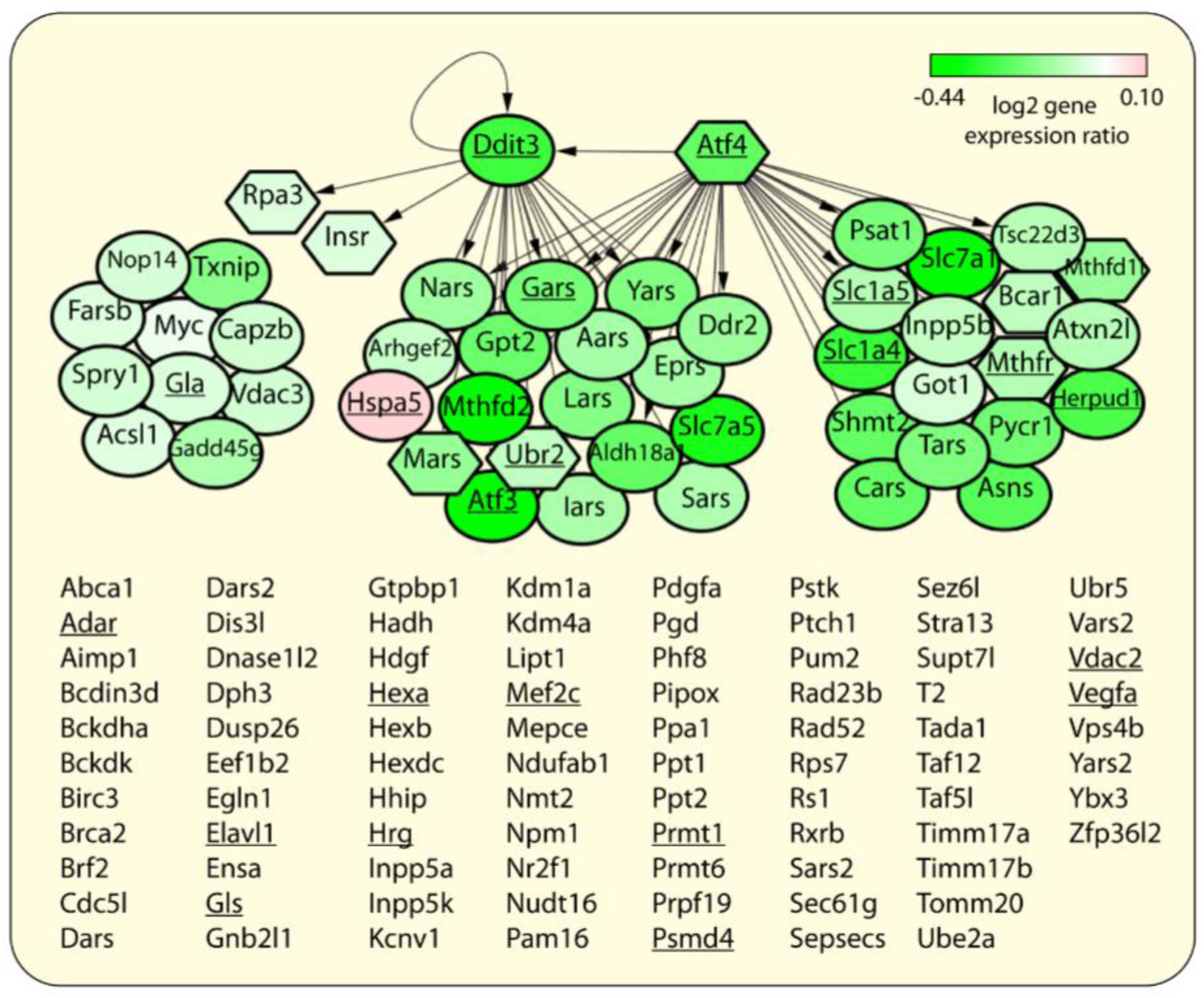
The 134 genes from the ALS gene expression classifier. Genes that are either transcriptionally responsive to ER stress [32] or directly targeted by TFs ATF4 (Atf4) and CHOP (Ddit3) during ER stress [32] are shown as nodes (depicted using Cytoscape [33]). Node color indicates mean log2 differential expression level in the indicator cell assay (disease versus normal) ranging from –0.44 (green) to 0.1 (pink). Node shape indicates genes that are transcriptionally responsive to ER stress (oval) or are not (hexagon) [32]. 133 of the 134 genes are known to have human orthologs and underlined genes co-occur with "amyotrophic lateral sclerosis" in titles or abstracts of articles in PubMed (on 01/15/15).

Secondly, the genes in the classifiers are significantly enriched for those differentially expressed in microdissected neurons from pre-symptomatic *SOD1*-mutant mice versus control mice[7–9] (HGD p-Values = 1.18E-06, 2.75E-05, and 6.78E-05 for Classifiers 1–3, respectively), and ALS-related genes in the ALSoD database on 01/06/2015[34] (HGD p-Values = 5.18E-3, 8.77E-3, and 0.03, for Classifiers 1–3, respectively). The gene-set classifiers also contain ALS-associated processes including ‘stress granule assembly’[35] and ‘positive regulation of p38MAPK cascade’[36] (containing risk loci[35] ATXN2 and VEGFA (VEGF), respectively). Classifier 3 uniquely contains up-regulated gene sets implicated in ALS biology, including ‘synaptic vesicle priming’[35] (containing ALS risk locus UNC13A[35]), as well as ‘glutamine catabolic process’ and ‘activation of AMPA [glutamate] receptors’, which is likely related to glutamate excitotoxicity in ALS[35,37]. These data suggest potential use of iCAP to model disease pathophysiology and to identify potential drug targets.

It is noteworthy that while CHOP/ATF4 targets related to protein synthesis appear up-regulated in microdissected neurons of *SOD1* mutant mice (a network state promoting ER stress-induced cell death [32]), almost all of these genes were down-regulated in the indicator cell assay (Fig 3; S2 Table), a network state consistent with cell survival [32]. This similar network structure, but different activity, suggests temporal dynamics of disease progression; the response of unaffected indicator cells after acute exposure to disease signals in serum is different than the response of affected neurons that have been persistently exposed to disease signals *in vivo*. Perhaps analyses of time-course iCAP data will eventually also reveal dynamics of ALS progression to unaffected neurons.

### An iCAP assay for early detection of Alzhiemer’s disease from human plasma samples

To test the applicability of the iCAP to different diseases, and the complex genetic and environmental diversity of a human population, we initiated development of an iCAP to detect AD at preclinical and early-symptomatic stages using a limited number of human plasma samples. For this application, we analyzed samples from patients with preclinical and early symptomatic stage AD, and normal healthy controls (defined in Table 3), with the assumption that the preclinical patients have the pathophysiology of AD, i.e. with amyloid deposition in brain that is indicated by a measured decrease of CSF abeta42 concentration, while not yet presenting clinical symptoms. Indicator cells for this experiment were a commercially available, pan-neuronal population of glutamatergic and GABAergic neurons derived from induced pluripotent stem cells (iPSCs) (from Cellular Dynamics International (CDI)), chosen because glutamatergic neurons are selectively degraded in AD [38] by extrinsic signals of disease [39]. Neurons were exposed to plasma samples for 24 h and RNA was isolated and used for gene expression analysis with Affymetrix human exon microarrays (ST 1.0). The data for all samples were merged, normalized [11], and analyzed for significant differential expression of genes and gene sets [13] between normal (N=20) and disease classes (N=38).

These data were used to generate AD classifiers using genes and clusters of co-regulated genes as features. Candidate genes were selected based on a combined score of magnitude of differential expression (fold change ratio), significance of differential expression (t-test p-Value), and significance of differential gene set expression (using GSA algorithm [13]). Candidate clusters of co-regulated genes (across all samples) were selected using clustering software [19], and filtered by feature selection [14]. Genes and gene clusters were used to train an SVM-based disease classifier. Performance was evaluated using leave-one-out cross-validation where all feature selection steps utilizing disease status occur inside the loop. This classifier achieved 72% accuracy (MCC p-Value = 3.10E-03) with an AUC of 0.64 (p-Value = 2.90E-2) (Table 4) and performed similarly with preclinical and early symptomatic AD test samples (data not shown). This preliminary analysis with unoptimized assay parameters, is presented to demonstrate proof-of-concept that the assay will work with human samples. However, the performance of the classifier is not yet optimal. Because the assay uses cells as signal transducers, numerous experimental parameters can be refined, and because of the variation associated with human samples, more samples need to be included to develop a highly accurate classifier. Future studies will include analyses with increased sample numbers, optimization of assay parameters, including plasma concentration and incubation time, and characterization and control of experimental variance.

As a first step towards improving classifier performance, we tested the effect of including a classifier feature based on patient *APOE* genotype, the strongest genetic risk factor for sporadic AD [40]. Including a count of the *APOE4* alleles for each patient as a feature of the iCAP classifier, improved the area under ROC curve over that achieved with the initial iCAP classifier (AUC = 0.69, p-Value = 4.68E-3). By comparison, a classifier based on only the *APOE4* count did not perform well (Table 4, S1 Fig). To assess classifier robustness, we created 100 16-fold cross-validated sets of predictions for each of the three classifiers. In these experiments, iCAP + *APOE4* performed better than iCAP or *APOE4* alone as measured by accuracy (p-Value < = 2.4E-11), MCC (p-Value < = 2.3E-2), and AUC (p-Value < = 2.1E-17) (based on the Wilcoxon signed rank test).

To investigate the biological significance of the cellular response, we analyzed the genes selected in these 100 16-fold cross-validation runs (S5 Table). The 128 genes selected in at least 50% of classifier iterations were used for functional enrichment analysis [41]. The top enriched disease annotation was ‘cognition disorders’ (FDR [18] = 1.46E-2), assigned to AD-linked genes *APOE, DRD2, DRD3* and *CRH* in the 128-gene list, and the top enriched pathway was G-protein coupled receptor (GPCR) ligand binding (FDR [18] = 1.10E-19) implicated in AD pathology [42], suggesting utility of the assay for understanding disease process and for drug discovery. Importantly, none of these 128 genes has a mouse ortholog in the 134 ALS gene set, implying that this signal is specific to AD and not part of a general neuron stress signature. This suggests that given sufficient training data, iCAP will provide an effective blood-based diagnostic tool for human diseases.

## Discussion

Low-cost, non-invasive blood-based assays are needed for many complex diseases, for early diagnosis and prognosis, monitoring response to treatment, and for developing therapies. It is recognized that such assays will likely involve multicomponent biomarkers that reflect multiple physiological axes.

We have established proof-of-concept for a platform for blood-based diagnostics that uses standardized cultured cells as biosensors that integrate disease signals in blood from potentially multiple physiological pathways involving multiple molecule types, and transforms them into an easily measurable multicomponent gene-expression output. The concept is founded on the idea that while direct and global analysis of blood for biomarker discovery is an emerging science, analysis of transcriptional responses of cultured cells to stimuli is well established. The platform can provide easily measured, multicomponent disease biomarkers from serum that could be standardized to clinical assessment scores for early detection, patient stratification, and analysis of disease progression.
iCAP’s broad utility was tested by developing assays for two different diseases. We developed an iCAP to accurately distinguish pre-symptomatic mice of an ALS mouse model from non-carrier littermates and from mice of a Huntington’s model using frozen serum samples. Challenging the assay to distinguish patients with pre- and early symptomatic AD from normal subjects using human plasma samples established proof of concept in a much more genetically and environmentally diverse human population. This work establishes a paradigm for capitalizing on the natural ability of cultured cells to detect and respond to complex disease signatures for multicomponent blood-based diagnostics with broad applicability to multiple diseases. As the iCAP takes an orthogonal approach to biomarker discovery, combining genetic or other biomarker data with iCAP data can lead to improved diagnostic accuracy [43], as we have shown for the AD iCAP.

We sought to compare iCAP’s performance to those of other assays under development. A blood-based ALS classifier that uses levels of neurofilament light chain protein (NfL) [44,45] achieved 71–75% specificity and 89–90% sensitivity at differentiating patients diagnosed with ALS from healthy controls [44]. There are also CSF biomarker assays in development; one such assay has independently validated 83% sensitivity and 100% specificity of identifying patients diagnosed with ALS versus patients without ALS [46]. Promising studies have also identified protein level changes in fibroblasts of ALS patients [47] and properties of erythrocytes [48] that could be good biomarkers, but classifiers have not yet been reported. It is feasible that similar markers could be derived from direct analysis of other cells isolated from patient blood, such as peripheral blood lymphocytes, which have shown promising signals relevant to ALS pathogenesis, suggesting systemic aspects of ALS that are detectable in blood [49]. However, because of noise due to the variable abundance of particular cell types in blood, genetic variation between individuals [50], and the prominent responses of circulating immune cells to generic proinflammatory signals [51], direct analyses of blood cells has inherent challenges for biomarker discovery. Supporting this, an approach using microarray signatures of blood cells could distinguish patients with a variety of autoimmune diseases from normal controls, but could not distinguish patients with autoimmune diseases from each other (including those with multiple sclerosis, systemic lupus erythematous, rheumatoid arthritis and insulin dependent diabetes mellitus) [52].

In contrast, the iCAP readout is from genetically identical and reproducibly obtainable, standardized indicator cells derived from stem cells, which can be chosen to be responsive to extrinsic signals of disease (e.g. dopaminergic neurons for Parkinson’s disease [53] or normal large airway epithelial cells for lung cancer [54]). The ALS feasibility study used cells in EBs that were differentiated in-house, but various human iPSC-derived cell types (approaching good manufacturing practices and International Organization for Standardization standards) are now commercially available, such as the iCell^®^ Neurons used for the AD assay.

Clinical application of the iCAP for early diagnosis of ALS, AD or other diseases faces complex variation in underlying genetic and environmental factors amongst individuals. The AD study tested robustness of the assay to this variation in a human population, suggesting that despite complex diversity amongst individuals, there are pathobiological similarities amongst the AD patients that can be detected by the assay. The variation tested in the ALS study was limited to the underlying genetic diversity of the transgenic mice [55], which likely contributes to the variability of disease onset of the model [5]. Nonetheless, we expect it is possible to develop a robust iCAP for clinical diagnosis of ALS because despite genetic heterogeneity underlying ALS, patients appear to have common underlying pathogenesis. For example, although mutant *SOD1* is the pathological cause of only 20% of familial ALS patients, *SOD1* is also implicated in sporadic ALS comprising 90% of all ALS cases [56]. However, the impetus of capturing multicomponent signatures is to not only identify general aspects of a disease, but to also find elements that stratify patients with the same disease for precision medicine. We look forward to exploring this possibility with larger human cohorts in the future.

An effective clinical iCAP would also need specificity to differentiate disease mimics with similar clinical symptoms (such as ALS and spinal muscular atrophy), and diseases with similar underlying molecular networks (such as ALS and Huntington’s disease). This capacity is supported by recent work demonstrating that diseases with phenotypic or pathobiological similarities have overlapping but not identical molecular networks [57], and by our data showing specificity of the murine ALS assay to ALS and not Huntington’s disease and the lack of overlap between AD and ALS classifier genes. For generating a clinical iCAP, disease mimics would be included in classifier training to improve selection of disease-specific responses.

Clinical deployment of iCAP assays would also require technical modifications for cost-effectiveness. Transitioning from using microarrays for gene expression analysis to using a targeted RNA detection platform is possible because deployment of the assay would require analysis of only the small subset of genes that inform the classifier along with appropriate controls. A second possibility is transitioning to another readout of the cellular response such as detection of fluorescent reporter proteins or metabolic profiling. Optimization of the readout and other optimizations, such as standardization, automation, and miniaturization of the growth and treatment of the indicator cells, provide a path to broad, robust and cost-effective research and clinical applications of the iCAP.

There are limitations to the iCAP approach. Blood analytes are not directly identified by the assay and can only be predicted through network modeling of the cellular response signature. In addition, the assay is more complicated than direct detection assays and involves cellular biosensors, the performance of which may be sensitive to batch and well variability and culture conditions. For these reasons, the effects of these parameters must be carefully monitored and standardized with appropriate controls, and clinical applications might best be performed in a single central performance site.

For the ALS and AD studies, the indicator cell types were selected based on their known involvement in the disease pathologies, but we do not yet know the limitations on type and species of origin of indicator cells. We expect that for some iCAP applications, a single cell type could be identified that can indicate multiple diseases. Success of the iCAP does not depend on understanding the cellular responses; at first blush, the response pattern is irrelevant as long as the classifier is robust. Nevertheless, the assay captures cellular responses to complex signals of active disease generated *in vivo*, and thus has unique relevance to disease pathobiology compared with many other *in vitro* models. This is particularly relevant for cancers [4] and neurodegenerative diseases [3,53], for which pathologies spread to unaffected cells via secreted material. Inspired by the presence of known disease-related elements in our data and the clinical need for understanding disease processes in inaccessible tissues, generating patient-specific *in vitro* models of disease from frozen serum samples is an appealing future application for the iCAP.

## Acknowledgements

We thank Dr. Michael Weiss for helpful discussions and suggestions. We are grateful to Drs. Thomas Jessell and Hynek Wichterle for providing the HBG3 embryonic stem cell line. We are grateful to Curis, Inc. for providing the hedgehog agonist, Hh-Ag1.3, used for stem cell differentiation.

## Supporting information

S1 Fig. ROC curves of AD classifier performances

S1 Table. 64 down-regulated gene sets used for iCAP-based Disease Classifier 1: Gene Set Classifier

S2 Table. 134 genes used for iCAP-based Disease Classifier 2: Gene Expression Classifier

S3 Table. 106 up- and down-regulated gene sets used for iCAP-based Disease Classifier 3: Gene Set Classifier

S4 Table. Classification of Huntington’s disease samples by the ALS classifier using 16 different classifier configurations

S5 Table. Ranked list of AD classifier genes selected at least once in the 100 16-fold cross-validation runs

